# Bellymount enables longitudinal, intravital imaging of abdominal organs and the gut microbiota in adult *Drosophila*

**DOI:** 10.1101/741991

**Authors:** Leslie Ann Jaramillo Koyama, Andrés Aranda-Díaz, Yu-Han Su, Shruthi Balachandra, Judy Lisette Martin, William B. Ludington, Kerwyn Casey Huang, Lucy Erin O’Brien

**Affiliations:** Department of Molecular and Cellular Physiology, Stanford University School of Medicine, Stanford, CA 94305; Department of Developmental Biology, Stanford University School of Medicine, Stanford, CA 94305; Department of Bioengineering, Stanford University, Stanford, CA 94305; Department of Embryology, Carnegie Institution of Washington, Baltimore, MD 21218; Department of Microbiology and Immunology, Stanford University School of Medicine, Stanford, CA 94305; Chan Zuckerberg Biohub, San Francisco, CA 94158

## Abstract

Cell- and tissue-level processes often occur across days or weeks, but few imaging methods can capture such long timescales. Here we describe Bellymount, a simple, non-invasive method for longitudinal imaging of the *Drosophila* abdomen at sub-cellular resolution. Bellymounted flies remain live and intact, so the same individual can be imaged serially to yield vivid time series of multi-day processes. This feature opens the door to longitudinal studies of *Drosophila* internal organs in their native context. Exploiting Bellymount’s capabilities, we track intestinal stem cell lineages and gut microbial colonization in single flies, revealing spatiotemporal dynamics undetectable by previously available methods.

---

A major thrust of modern biology is leveraging advancements in live microscopy to reveal how cellular and physiological processes unfold inside living organisms. For adult metazoans, achieving this goal requires overcoming two imaging challenges: the opacity of most mature animals to light, and the prolonged timescales of adult-associated processes such as ageing.

The adult vinegar fly, *Drosophila melanogaster*, has yielded foundational insights into metazoan physiology^1–4^. This invertebrate animal is also a powerful tool for probing human disease pathologies, with ∼65% of human disease-causing genes having functional homologs in the fly^5^. Current methods for imaging *Drosophila* abdominal organs, however, are limited in optical resolution, imaging duration, or both. Some newer approaches preserve animal viability but cannot visualize individual cells^6–8^. Other recent advances enable high-resolution imaging but require opening the abdominal cuticle, which leads to death of the fly^9–12^.

Here we present Bellymount, a method for high-resolution imaging of the intact *Drosophila* abdomen. Bellymount captures volumetric images of native abdominal organs at spatial scales ranging from subcellular (<1 µm) to multi-organ (>100 µm). It preserves organismal viability, thus enabling for the first time longitudinal studies of *Drosophila* abdominal organs and cells. It is inexpensive to construct, simple to apply, and compatible with diverse brightfield and fluorescence microscopes. Finally, Bellymount is easily combined with *Drosophila*’s sophisticated tools for spatiotemporal genetic manipulation, fluorescent labeling, and live reporter assays. Exploiting all these features, we use Bellymount to perform longitudinal tracking of two multi-day processes in the fly gastrointestinal tract, generation of intestinal stem cell lineages, and colonization of the gut by commensal bacteria. The resulting time series provide the first direct views of spatial and temporal heterogeneities that underlie both events, and demonstrate the capability of Bellymount to uncover new physiological dynamics of cells, tissues, and organs *in vivo*.

The exterior cuticle of the adult fly, which is generally opaque, presents an obstacle for light-based imaging of internal organs. Serendipitously, we noticed that the ventral abdominal cuticle becomes transparent when affixed to a glass coverslip by the polyvinyl acetate adhesive, Elmer’s Clear School Glue (Fig. 1a). We named this procedure ‘Bellymount’. Transparency of the glued cuticle enabled facile observation of organs such as the midgut, crop, and female ovaries (Fig. 1c) in flies that were live and intact. Bellymounted flies were readily removed from the coverslip, even hours after the glue had dried (Supplemental Movie 1), and they typically remained viable. In a survival assay, 92% of flies were alive 24 hours after being glued and released (Supplemental Fig. 1).

**Figure 1:**
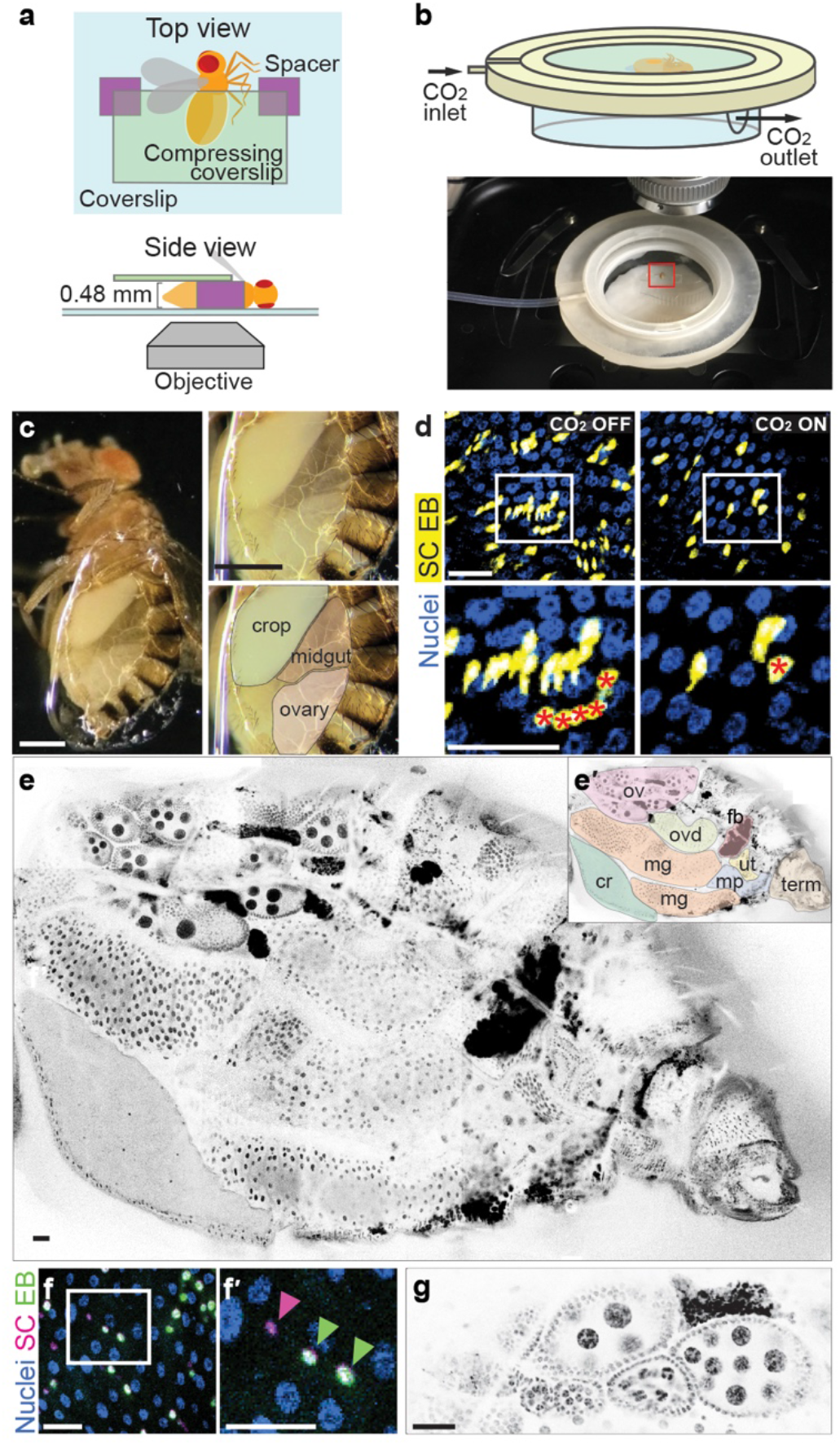
Bellymount enables intravital imaging of abdominal organs in live adult *Drosophila* at micron-resolution. a, Cartoon of Bellymounted fly. The cuticle of the ventrolateral abdomen is glued to the coverslip. To maximize contact with the glue, gentle pressure is applied to the top of the abdomen with a second, compressing coverslip atop two spacers. b, Bellymount apparatus. *Top*, Schematic of 3D-printed apparatus. The coverslip with the glued fly is nested inside. CO2 flows through the apparatus via the indicated ports. *Bottom*, Bellymount apparatus positioned for imaging on upright microscope. Red box shows the position of the glued fly. See Supplemental Fig. 2. c, Gluing causes the ventral cuticle to become light-transparent. *Left*, Adult female with ventrolateral abdomen glued and compressed as in Fig. 1a. The edge of the glue patch appears as a refractive oval line around the abdomen. *Right*, The crop, the midgut, and an ovary are visible through the glued cuticle. See Supplemental Movie 2. d, Carbon dioxide (CO2) minimizes tissue movements for confocal fluorescence microscopy. *Left*, Without CO2, tissue movements during *z*-stack acquisition resulted in multiple representations of the same cells. *Right*, With CO2, movements were inhibited, and no multiple representations are present. In zoomed panels (bottom), red asterisks label the identical cell in images taken without and with CO2. Nuclei (His2av::mRFP) are shown in blue; midgut stem cells and enteroblasts in yellow (LifeactGFP). See Supplemental Movie 3. e-g, Intravital, micron-resolution imaging of multiple abdominal organs. e, Projection of a tiled z-stack reveals arrangement of abdominal organs in a live, intact female. Nuclei (His2av::mRFP) in inverted grayscale. e’, Visible organs are: crop (cr), mg (midgut), ov (ovary), ovd (oviduct), fb (fat body), ut (uterus), mp (Malphigian tubules), term (terminalia). f, Multichannel view of cell types in the midgut. Stem cells (SC, red nuclei) and immature enteroblasts (EB, green-yellow nuclei) were dispersed among mature enterocytes (large blue nuclei). f’, Zoomed region with nuclei of stem cell (pink arrowhead) and two enteroblasts (green arrowheads). g, Detail of ovary shows egg chambers at different developmental stages. Nuclei (His2av::mRFP) in inverted grayscale. Panels d-g are projections of confocal stacks. Genotype of flies in d: *esg>LifeActGFP*; *ubi-his2av::mRFP*. Genotype of flies in e-g: *esg>his2b::CFP, GBE-Su(H)-GFP:nls*; *ubi-his2av::mRFP*. All scale bars, 30 µm.

The abdomen is the body’s central location for digestive physiology and function. To explore the applicability of Bellymount for gastrointestinal studies, we used an inexpensive, USB-pluggable microscope to record food ingestion and transit (Supplemental Movie 2). Bellymounted flies were provided 5% sucrose water that was colored with Brilliant Blue FCF. Over 45 min, the ingested blue liquid filled successive compartments of the gastrointestinal tract, with rapid peristaltic contractions accompanying nutrient transit. The sharp resolution of these digestive events in time and space demonstrates the potential of Bellymount to investigate gastrointestinal function in real time.

We next assessed whether we could resolve cells in the midgut using confocal microscopy. Initially, midgut peristalsis and global body movement during image acquisition caused fluorescently labeled cells to appear duplicated (Fig. 1d, Supplemental Movie 3). To overcome this issue, we designed a custom apparatus to apply anesthesia via carbon dioxide (CO_2_) exposure. The Bellymount apparatus comprises three parts: a base to hold the coverslip with the glued fly, a screw-on lid, and a humidity chamber to ensure that the fly does not desiccate during imaging (Fig. 1b, Supplemental Fig. 2). The base and lid were 3D-printed at minimal cost (Methods). CO_2_, delivered through an inlet in the base, inhibited both midgut peristalsis and overall body movement. This effect enabled acquisition of crisp images at subcellular resolution (Fig. 1d, Supplemental Movie 3).

**Figure 2:**
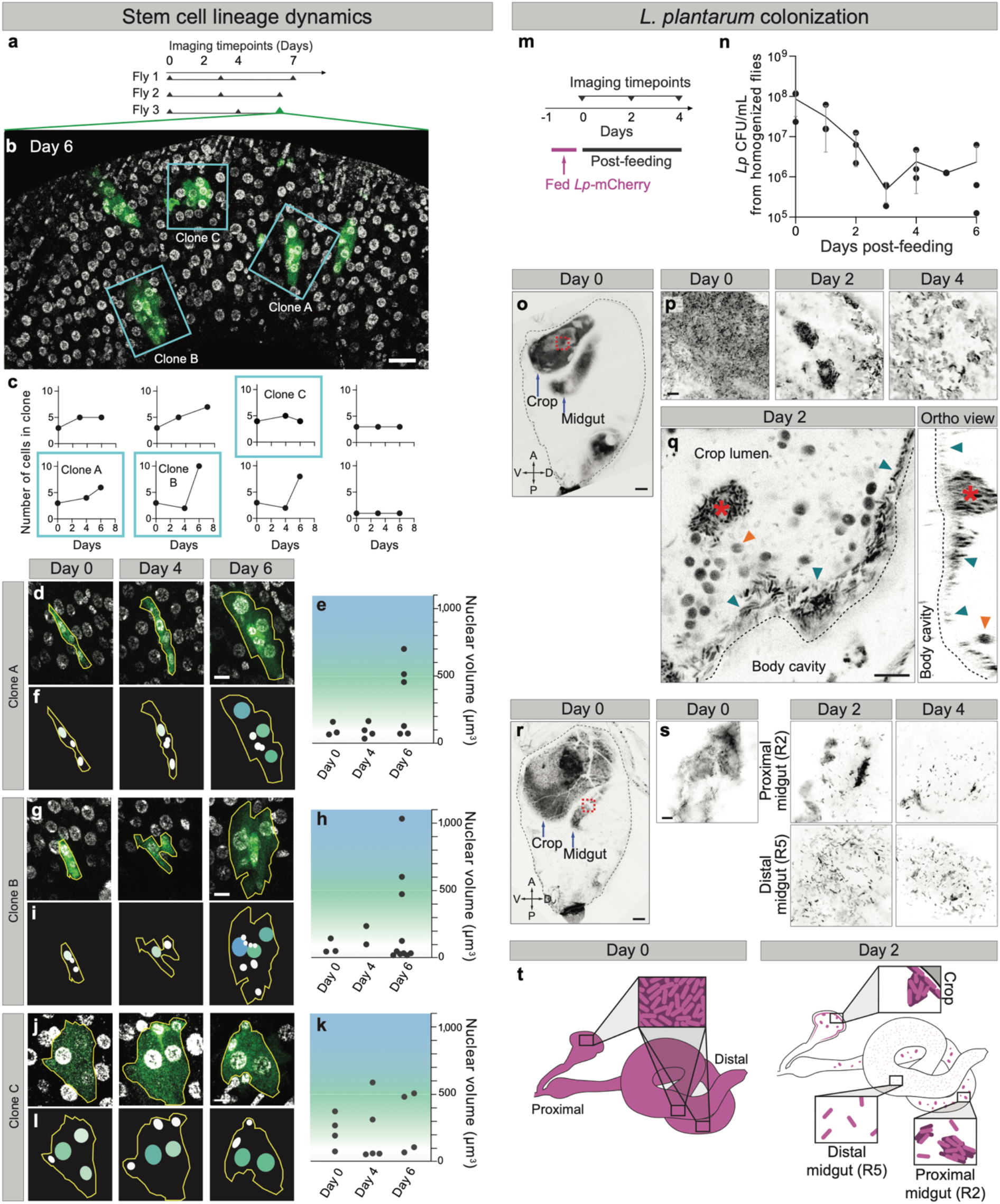
Serial Bellymount imaging reveals longitudinal dynamics of midgut stem cells and gut bacteria. a-j, Dynamics of midgut stem cell lineages: a, Timepoints for serial imaging of GFP-marked stem cell clones. Clones arose spontaneously in the first 4 days of adult life and were imaged 3 times over the subsequent 6-7 days. Day 0 designates the first imaging session. b, Clones formed organ-level patterns that enable their re-identification across imaging sessions. Blue boxes outline the midgut clones in Fly 3 that were tracked for the duration of the experiment. Green, GFP-labeled clones. Grayscale, nuclei (His2av::mRFP). Scale bar, 30 µm. See Supplemental Fig. 4. c, Longitudinal dynamics of clone cell addition and loss. Cells were counted at each imaging timepoint for 8 clones from the 3 flies in (a). d, g, j, Zoomed images of the tracked midgut clones in (b) at each timepoint. Scale bar, 10 µm. e, h, k, Longitudinal dynamics of clone cell differentiation as revealed by nuclear volume. Larger nuclear volumes (green to blue background) represented more advanced stages of enteroblast-enterocyte differentiation, while stem cells had small nuclear volumes (white background). Small nuclei are also a feature of enteroendocrine cells, which cannot be distinguished from stem cells in this experiment. See Supplemental Fig. 6. f, i, l, Cartoons of clones in (d, g, j). Colors of nuclei in cartoons correspond to the colored background for nuclear volumes plotted in (e, h, k). m-t, Gut colonization by *L. plantarum*: m, Time points for serial imaging of *L. plantarum* colonization. Flies were provided a 1-day pulse of *L. plantarum*-mCherry. After *L. plantarum* removal, individual flies were imaged 3 times over 4 days. Day 0 designates the first imaging session. n, Colony forming units (CFUs) of *L. plantarum* over time. A cohort of flies was homogenized at the indicated time points after the 1-day pulse of *L plantarum*-mCherry. Levels of *L*. *plantarum* were high immediately after the pulse, decreased over the next 3 days, and then plateaued, indicating stable colonization. Each point represents CFUs from one fly. o-q, Longitudinal tracking of *L. plantarum* colonization in the crop. o, Single optical section shows whole-abdomen tilescan at day 0 after feeding *L. plantarum*-mCherry (inverted greyscale). Red box indicates Day 0 zoomed-in region in (p). Scale bar, 30 µm. p, Single optical sections show spatial patterns of *L. plantarum*-mCherry in the same crop on days 0, 2, 4. Inverted greyscale, *L. plantarum*. q, High-magnification optical section (*left*) and orthoview (*right*) of crop from (p), Day 2. Bacteria accumulated at the edge of the crop (blue arrows) and formed prominent clumps (red asterisk). Ingested yeast cells (orange arrows) were also visible due to autofluorescence. See Supplemental Movie 6. r-s, Longitudinal tracking of *L. plantarum* colonization in the midgut. r, Single optical section shows whole-abdomen tilescan at day 0 after feeding *L. plantarum*-mCherry (inverted greyscale). Red box indicates Day 0 zoom region in (s). Scale bar, 30 µm. s, Single optical sections show spatial patterns of *L. plantarum*-mCherry in the same midgut on days 0, 2, 4. On day 0 only one region of the midgut was visible. On days 2, 4 two regions of the midgut became visible. Distinct spatial patterns of bacterial colonization characterized proximal and distal midgut regions. Inverted greyscale, *L. plantarum*. See Supplemental Movies 7, 8. t, Heterogenous dynamics of bacterial colonization and dispersal in distinct regions of the GI tract as revealed by longitudinal tracking. Individual regions of the GI tract including the crop, proximal midgut, and distal midgut exhibited distinct spatial patterns of bacterial localization that developed within an individual over time. Genotype for (b-l): *UAS-CD8-GFP, hs-flp; tubGal4; FRT82, tubGal80/ubi-his2av::mRFP, FRT82*. Genotype for (m-t): *ubi-his2avD::YFP.* Unless indicated, images are projections of confocal stacks. All scale bars, 10 µm unless otherwise indicated.

By gently compressing the abdomen of the glued fly during anesthesia (Fig. 1a), we were able to view the whole abdomen at single-cell resolution. Using either confocal or two-photon microscopy, we acquired volumetric tile scans that included portions of nearly all female abdominal organs: midgut, crop, rectum, ovary, oviduct, uterus, trachea, fat body, and Malpighian tubule, as well as circulating hemocytes (Fig. 1e, Supplemental Fig. 3, and Supplemental Movie 4). In the midgut, stem cells and various stages of terminal progeny were easily distinguished when labeled with fate-specific fluorescent markers (Fig. 1f). In the ovary, egg chambers at various developmental stages were apparent; within these chambers, nascent oocytes and their supporting cells were readily identifiable (Fig. 1g and Supplemental Movie 5). Thus, Bellymount enables—for the first time in adult *Drosophila*—observation of native abdominal organs at single-cell resolution, in animals that are live and intact.

The viability of flies after Bellymount raised the possibility of performing serial Bellymount imaging on the same individuals for longitudinal studies. To investigate this possibility, we evaluated the suitability of Bellymount for tracking cellular events in single flies over multiple days. We focused on two types of cellular events, (1) divisions of midgut intestinal stem cells (Fig. 2a-l), and (2) microbial colonization of the gastrointestinal tract (Fig. 2m-t). Both processes are conserved in vertebrates and have attracted intense research interest, yet their real-time dynamics remain virtually unexplored due to lack of methods for multi-day tracking in single flies.

The *Drosophila* midgut is physiologically equivalent to the vertebrate stomach and small intestine. As in vertebrate intestine, stem cells in the fly midgut continuously divide to replenish terminally differentiated epithelial cells that form the intestinal barrier^13^. Although current live imaging methods capture single divisions^12^, their timescales are not sufficient to track multiple divisions of the same stem cell or to monitor differentiation of cellular progeny.

To reveal a stem cell’s division history, generating marked clones is the gold standard^14^. Constitutive expression of a marker such as GFP is enabled in a small number of single stem cells, and progeny that arise from these labeled stem cells inherit marker expression, enabling their identification. Over time, these progeny manifest as a cluster of labeled cells, termed a clone, and the cellular composition of the clone represents the stem cell’s ‘lineage’.

Midgut stem cell clones traditionally have been analyzed in fixed samples. This approach yields a static snapshot of a stem cell’s lineage. However, it does not provide the temporal history of when progeny were born, how quickly they differentiated, or whether any progeny died. Such time-resolved information is crucial for a deep understanding of adult tissue homeostasis.

We asked whether the histories of single stem cell lineages could be tracked by serial imaging using Bellymount. Stem cell lineages were labeled using MARCM (Mosaic Analysis with a Repressible Cell Marker)^15^. Spontaneous recombination during the first 4 days of adult life generated sporadic GFP-marked midgut stem cells, which subsequently developed into GFP-marked, multicellular clones (Fig. 2b and Supplemental Fig. 4).

To test whether specific clones could be re-identified over time, we serially imaged 3 flies on 3 occasions over 6-7 days (Fig. 2a). In these individuals, 8 clones were re-identifiable because they exhibited unique spatial patterns or distinctive shapes (Supplemental Fig. 4). Other clones could not be tracked because they lacked distinguishing morphological characteristics or disappeared from view due to slight displacements or rolling of the midgut tube.

We investigated the longitudinal dynamics of these 8 stem cell lineages. First, we determined how lineages grew or shrank by counting the number of clone cells per timepoint (Fig. 2c). This analysis revealed diverse trajectories: 2 clones kept the same number of cells, 3 clones added additional cells, and 3 clones not only added cells but also lost them—events that could not have been detected in fixed tissues. Between timepoints, rates of cell addition exhibited a 16-fold range of 0.25-4 cells per day. These heterogenous dynamics carry implications for how stem cell lineages evolve. While numbers of cells in our tracked clones resemble those reported in fixed tissue studies^16–18^, our longitudinal analysis with Bellymount revealed that clones with similar numbers of cells can arise through highly distinct trajectories.

Next, we considered the rates at which cells differentiated into enterocytes. In the fly midgut, a newborn stem cell daughter differentiates into a terminal enterocyte via an intermediate state called an enteroblast. This process is characterized by increasing ploidy; stem cells and new daughters are 2N, enteroblasts are 2-8N, and enterocytes are 8-64N^18^. Hence, nuclear volume provides an indicator of how far immature enterocytes have progressed toward terminal differentiation (Supplemental Fig. 5).

Using nuclear volumes, we assessed differentiation rate for cells in three clones from one midgut (Clones A, B, and C in Fly 3; shown in organ-level view in Fig. 2b and Supplemental Fig. 5 and in zoomed view in Fig. 2d,g,j). These measurements indicated that some stem cell progeny progressed to enteroblast or enterocyte states whereas others did not (Fig. 2e,f,h,i,k,l). In specific cases, the data also provided more nuanced information: In Clones A and B, enterocytes were absent between days 0-4 but present at day 6, suggesting that, in some cases, enterocyte differentiation proceeded rapidly once initiated. In Clone C, the disappearance of an enterocyte between days 0 and 4 implied either cell loss or de-differentiation.

Altogether, these clonal analyses provide the first direct views of how single stem cell lineages develop. They show that dynamics of new cell addition and differentiation vary widely, not only between different stem cell lineages^17, 19^, but also within the same lineage at different times. The ability of Bellymount to track individual lineages for prolonged times will facilitate future mechanistic studies of these dynamics.

We applied serial Bellymount imaging to examine a second dynamic process, colonization of the gastrointestinal tract by commensal bacteria. While the human gut microbiome comprises hundreds of bacterial species, the *Drosophila* gut microbiome typically comprises only five^20^. This relative simplicity, together with *Drosophila*’s genetic tractability, has helped establish the fly as a powerful model for mechanistic study of host-microbiome interactions.

The biogeography, or dynamic spatial distribution, of the gut microbiota along the length of the gastrointestinal (GI) tract is known to impact host-microbiota interactions and digestive physiology^21, 22^. However, the biogeography of the *Drosophila* gut is largely mysterious because no existing methods can monitor gut bacteria throughout the GI tracts of living flies.

We examined whether serial Bellymount imaging could provide a direct view of gut microbial colonization. The prevalent and abundant gut commensal *Lactobacillus plantarum* was tagged with mCherry^23^ and fed to conventionally reared flies for 1 day. The next day, flies were removed from *L*. *plantarum*-mCherry and, for the remainder of the experiment, maintained on fresh food (Fig. 2m). Measurements of colony forming units in a cohort of homogenized flies confirmed that levels of *L. plantarum* were high immediately after the 1-day pulse, decreased over the next 3 days, and subsequently plateaued, indicating stable colonization^23^ (Fig. 2n).

We performed whole-abdomen volumetric imaging immediately (0 days), 2 days, and 4 days after the *L. plantarum* pulse (Fig. 2m, o, and r, respectively) to reveal the bacteria’s spatial distribution. At 0 days, *L. plantarum*-mCherry densely occupied the lumens of both the crop, a proximal storage organ^24^, and the midgut, with a filling fraction of 51±15% (*n*=9). In the same individuals 4 days later, *L. plantarum*-mCherry was sparse in both organs, with a filling fraction of 4.6±4.7% (*n*=5) (Fig. 2p,s). These observations were consistent with CFU measurements.

Beyond filling fraction, the three-dimensional patterns of *L. plantarum* exhibited intriguing regional and temporal changes that could not have been detected in homogenized flies. During colonization of the crop, *L. plantarum* localized to the crop wall, where it frequently coalesced into prominent clumps (Figs. 2q,t; Supplemental Movie 6). By contrast, in the midgut *L. plantarum* remained in the lumen, where it formed clumps in the proximal midgut while remaining dispersed as single cells in the distal midgut (Fig. 2s,t; Supplemental Movies 7, 8). These differences are consistent with impaired bacterial viability after transit through the acidic environment of the middle midgut^25^. Altogether, this time-resolved analysis of *L. plantarum* colonization provides the first insights into the dynamic regional biogeography of the *Drosophila* gut microbiota.

In summary, Bellymount enables longitudinal studies in the *Drosophila* abdomen through serial, micron-resolution imaging of flies that remain live and intact. Using Bellymount, we observed real-time digestive transit, visualized the native arrangement of abdominal organs, and resolved the organs’ constituent cells and resident microbiota. We applied serial Bellymount imaging to perform time-resolved tracking of midgut stem cell lineage dynamics and gut bacterial colonization, two multi-day processes that were previously inaccessible to live study. These experiments revealed previously undescribed heterogeneities in the spatial and temporal events that underlie midgut physiology and host-microbiome interactions. The Bellymount platform, which is based on Elmer’s Glue and a simple, 3D-printed apparatus, is inexpensive to implement, versatile in application, and compatible with a wide range of upright and inverted microscope systems. These features will facilitate the use of Bellymount to study the real-time dynamics of the diverse cellular and physiological processes that occur over prolonged timescales in adult animals.

## Methods

### Drosophila stocks

We obtained *hemolectinGal4; UAS-2xEGFP* (BL30140), *ubi-his2av::mRFP* (BL23650), and *10xUAS-IVS-myr-td::Eos* (*UAS-Eos*) (BL32226) from the Bloomington Stock Center. *esgGal4* (112304) was obtained from the Kyoto *Drosophila* Genetic Resource Center. The following stocks were gifts: *mexGal4* (Carl Thummel), *breathlessGal4*, *UAS-his2b::CFP* (Yoshihiro Inoue), *ubi-his2avD::YFP* (Pavel Tomancak), and *GBE-Su(H)-GFP:nls* (Joaquin de Navascues), *UAS-CD8-GFP, hs-flp*^12^*; tubGal4; FRT82, tubGal80* and *FRT82* (David Bilder). The ‘fate sensor’ line (*esgGal4, UAS-his2b::CFP, GBE-Su(H)-GFP:nls; ubi-his2av::mRFP*) was generated in a previous publication^12^.

### Drosophila husbandry

Flies and crosses were kept at 25 °C unless otherwise indicated. All experiments were performed on adult females. Flies were raised on standard cornmeal molasses media (water, molasses, cornmeal, agar, yeast, Tegosept, propionic acid). Unless otherwise indicated, following eclosion, flies were kept on cornmeal molasses vials supplemented with a pinch of powdered dry yeast (Red Star, Active Dry Yeast) with males for 4 days prior to imaging.

### Fabrication of Bellymount apparatus and humidity chamber

The Bellymount apparatus consists of a lid, base, and humidity chamber (Supplemental Fig. 2). The base and lid were 3D-printed using the online service Shapeways (https://shapeways.com). Fabrication was performed with fine detail plastic (Visijet M3 Crystal UV curable plastic) and the basic “smooth” finish option.

To prevent the fly from desiccating during imaging, we attached a humidity chamber to the underside of the apparatus base (Supplemental Fig. 2). Briefly, we drilled a 3-mm outlet for CO_2_ into the wall of a 35-mm petri dish (Olympus plastics, #32-103), then adhered the chamber to a small groove on the underside of the base using dental wax (Surgident, #50092189). Lastly, we covered the bottom of the Petri dish with trimmed paper towels moistened with H^2^O.

### Animal preparation

Flies were glued to the imaging coverslip (Fig. 1a) as follows: Flies were chilled on ice in a microfuge tube for at least 1 h before gluing to anesthetize them. Next, we painted a small rectangle of Clear Elmer’s School glue (Amazon, B06WVDBR62) roughly the size of the fly abdomen onto the center of a 40-mm coverslip (Fischer Scientific, #NC0018778) using a Worm Pick (Genesee, #59-AWP [handle] and #59-32P6 [tips]). After applying the glue, we quickly adhered flies to the coverslip.

Gluing the fly on its left ventrolateral surface provided optimal viewing of the gastrointestinal tract (Fig. 1c). To achieve the desired positioning, each of the fly’s two most posterior legs were held with a pair of Dumont #5 forceps while laying the fly’s ventrolateral side onto the Elmer’s glue. During gluing, care was taken to ensure none of the legs were trapped between the abdomen and the coverslip. After positioning the fly, we gently pressed the abdomen into the Elmer’s glue using a paintbrush.

Maximizing contact with the coverslip maximized visibility of abdominal organs. Therefore, we gently compressed the fly after gluing by placing a second, compressing coverslip atop the fly (Fig. 1a). The compressing coverslip was a square coverslip (Fischer Scientific, #12-541B) that had been broken in half. We found that 0.48 mm was the optimal distance between the compressing coverslip and the primary coverslip to ensure that the fly experienced compression without undue force. To position the compressing coverslip, we placed two 0.48 mm-thick, adherent spacers (Millipore-Sigma, #GBL620004-1EA) on either side of the fly and placed the compressing coverslip on top of the spacers. After thus securing the fly, we nested the coverslip inside the base of the apparatus and screwed on the lid.

### Anesthesia during Bellymount imaging

To minimize voluntary and involuntary tissue movements, we applied CO_2_ anesthesia during imaging. We delivered CO_2_ using 2-mm inner diameter (ID) flexible, silicone tubing attached to the inlet built into the base of the Bellymount apparatus (Fig. 1e, Supplemental Fig. 2). To prevent desiccation of the fly, the tubing was connected to a 500-ml Pecon humidification bottle containing distilled water. The humidified CO_2_ was piped through a secondary regulator (Micromatic, #8011-15), which allowed fine control of CO_2_ flow during imaging. The secondary regulator was attached, in turn, to a primary regulator and CO_2_ tank.

### Release of Bellymounted flies

To release flies from the Bellymount apparatus after imaging, the compressing coverslip and spacers were removed as a single unit and saved for future experiments. The tip of a pair of Dumont #5 forceps was placed under the fly thorax to gently pry the fly from the dried Elmer’s glue (Supplemental Movie 1).

Occasionally, a layer of Elmer’s glue remained on the fly’s abdomen after it was removed from the Bellymount apparatus. This layer could be peeled off easily by grabbing a free ‘tab’ of dried glue with a pair of forceps. If no free surface of dry glue was present, the glue was rewetted with milliQ water using a small paintbrush. After allowing the re-wetted glue to dry, a free tab of Elmer’s glue would commonly present itself. The glue was then peeled off with forceps as described above.

### Survival assay

We measured the lifespans of flies glued to a coverslip and compressed (see **Animal preparation**), to test for potential effects of the Bellymount protocol on viability. Flies were collected after eclosion and placed in molasses vials with powdered dry yeast and males for four days before the experiment. Flies were then randomly split into two groups (Bellymounted, *n*=50; control, *n*=49). Bellymounted flies were glued and compressed as described above (see **Animal preparation**), except that flies were held at room temperature in an empty pipette tip box with moistened paper towels rather than in the imaging apparatus. After 1 h, flies were released from Bellymount as described above (see **Release of Bellymounted flies**) and placed in a vial with 8-10 other experimental flies and 3-4 males. The control group did not undergo any Bellymount procedure and were maintained similarly as a control. We recorded the number of deceased flies each day and flipped the remaining survivors onto fresh food. Flies were maintained at 25 °C over the 7-day course of the experiment.

### Microscopy

We collected confocal images using four microscope systems: (1) an inverted Zeiss LSM880 with Zen software and Airy scan mode (Fig. 2m,n and Supplemental Figs. 3a, 6); (2) an inverted Zeiss LSM780 microscope with Zen software (Figs. 1h, 2o-q; Supplemental Movies 5-8); (3) an upright Leica SP5 confocal (Figs. 1b,c,f,g, 2c,d,f,h; Supplemental Figs. 3b, 4; Supplemental Movies 3, 4); and (4) an inverted Leica SP8 confocal (Supplemental Fig. 5).

We acquired brightfield images using three microscope setups: (1) a Zeiss Discovery V8 stereodissection microscope coupled with an iPhone 5S camera (Fig. 1a); (2) a Leica stereodissection microscope coupled with an iPhone X (Supplemental Movie 1); and (3) a digital, USB-pluggable microscope (Plugable.com, USB2-MICRO-250X Digital Microscope) (Supplemental Movie 2).

### Determining visible regions of the midgut

To determine which regions of the midgut are visualized by Bellymount imaging, we used Eos, a photoconvertible fluorophore^26^, to selectively mark visible regions of the midgut. Four day-old flies (*mexGal4, Gal80^ts^; ubi-his2av::mRFP / UAS-Eos*) were glued and prepared as described above (see **Animal preparation**). Using 405-nm laser light on a Zeiss LSM880 confocal, Eos protein was photoconverted from green to red emission.

After photoconversion, flies were removed from the Bellymount apparatus as described above (see **Release of Bellymounted flies**) and the midgut was examined *ex vivo* to determine the photoconverted regions. An 8-well Secure-Seal spacer sticker (ThermoFisher, #S24737) was used to form ‘wells’ on a microscope slide (Fischer Electronic Microscopy Sciences, #63720–05). After dissecting in Schneider’s Insect medium (Sigma-Aldrich, #S0146), one midgut and 7 µL of Schneider’s medium were placed in each well and topped with a coverslip (Fisher Scientific, #12-545-81). Midguts were imaged using an inverted Zeiss LSM 880 immediately after mounting.

### Time-lapse imaging of nutrient ingestion and midgut peristalsis

To monitor ingestion and midgut peristalsis, we performed low-magnification imaging of the fly abdomen while feeding dye. To easily identify the midgut for imaging, we fed FCF Brilliant Blue dye prior to the imaging experiment. Briefly, 24 h prior to imaging, flies were placed in a vial with blue-dyed yeast paste (10% FCF Brilliant Blue Dye (Sigma-Aldrich, #80717) in water mixed with dry yeast) and male flies.

After feeding for 1 day, flies were mounted as described above (see **Animal Preparation**). A cotton feeding wick was positioned in proximity to the fly’s proboscis and glued to the coverslip using KWIK-SIL silicone glue (World Precision Instruments, 60002). The feeding wick was attached to 2-mm ID flexible silicone tubing connected to a 10-mL syringe reservoir (Thermo Fischer Scientific, #03 377 23) filled with 10% FCF Brilliant Blue Dye in 5% sucrose (Sigma-Aldrich, #84097) water. Once the feeding wick had been properly placed and filled, the coverslip was gently placed into the apparatus with the tubing and reservoir attached through the CO_2_ outlet.

To image the fly, a USB pluggable microscope (Plugable.com, USB2-MICRO-250X Digital Microscope) was positioned above the abdomen of the fly and images were acquired every 0.5 s.

### Longitudinal imaging, tracking, and analysis of stem cell clones with MARCM

The MARCM system^15^ was used to generate GFP-marked stem cell clones.

MARCM turns on permanent, heritable GFP expression specifically in mitotic cells, when chromosomal recombination results in loss of GAL80^ts^ and consequent tubGAL4-driven expression of UAS-GFP in one daughter. In the adult fly midgut, MARCM specifically labels stem cell lineages because stem cells are, with rare exceptions, the only cells that undergo mitosis^16, 18, 27^.

Crosses for MARCM labeling were maintained at 18 °C prior to eclosion and shifted to 25 °C within the first 8 hours after eclosion. This temperature shift results in spontaneous GFP labeling of a small fraction of midgut stem cells. Four days after 25 °C temperature shift, we performed the first session of Bellymount imaging (day 0) (Fig. 2a). The sparseness of these spontaneous clones facilitated re-identification of clones during subsequent imaging sessions (Fig. 2a). After each imaging session, flies were placed in fresh vials (1 fly/vial) with powdered dry yeast and one male at 25 °C. Flies were flipped to new vials each day.

We identified and analyzed MARCM clones by examining serial confocal sections. Only clones that could be tracked for three consecutive time points following day 0 imaging were analyzed. We identified clones that were suitable for tracking by comparing the clone’s location relative to neighboring clones. Then, we mapped the clone arrangement to images of the same individual taken at other time points to verify that the clones were the same (Supplemental Fig. 4). In some cases, clones had a distinctive shape that allowed them to be identified across imaging sessions.

MARCM clone size and development were analyzed using FIJI and Bitplane Imaris v. 9.3.0. We inferred clone size by counting the number of contiguous cells in the discrete clone labeled with the nuclear marker *ubi-his2av::mRFP* and by summing the nuclear volumes of each cell within the clone. We computed nuclear volume using the Imaris contour tool by creating a surface from the *ubi-his2av::mRFP*-labeled nucleus and determining the enclosed volume.

### Preparation of fixed samples

Fixed samples with labeled cell types were used to determine the relationship between nuclear volume and differentiation state (Supplemental Fig. 5) as a baseline for analyses of differentiation rate (Fig. 2 e,f,h,i,k,l). Flies of genotype *esgGal4, UAS-his2b::CFP, GBE-Su(H)-GFP:nls; ubi-his2av::mRFP* were fixed, immunostained, and mounted as described previously^27^. Primary antibody: mouse anti-Prospero (1:400, DSHB, #MR1A), which stains enteroendocrine cells. Secondary antibody: Alexa Fluor 647-conjugated goat anti-mouse IgG (1:400, Thermo Fischer Scientific #A-21240). Samples were mounted in ProLong (LifeTechnologies). Stem cells, enteroblasts, and enterocytes were identified as described previously^12^. Nuclear volume for each cell type was assessed as described above (see **Longitudinal imaging, tracking, and analysis of MARCM clones**).

### *L. plantarum* colonization

For experiments with *L. plantarum*, flies were raised on standard cornmeal molasses food. Beginning with *L. plantarum* feeding and for the duration of the experiment, flies were shifted to cornmeal molasses food lacking Tegosept and supplemented with 10 µg/mL chloramphenicol (Calbiochem 220551).

A wild fly isolate of *L. plantarum* was tagged with a plasmid encoding mCherry and chloramphenicol resistance^23^. This strain was grown overnight from a frozen stock (25% glycerol) in 3 mL MRS medium (Difco^TM^ Lactobacilli MRS Broth, BD #288110) containing 10 µg/mL chloramphenicol. We spun down 1.5 mL of the saturated culture at 2000 rpm for 5 min at 4 °C. Next, we resuspended the pellet in 200 µL MRS + 10 µg/mL chloramphenicol and pipetted onto Whatman paper (Sigma-Aldrich, #WHA1002110) on fresh cornmeal molasses media as described above. Ten females and 3 males were transferred to each vial, which were wrapped in aluminum foil to prevent photobleaching of bacteria. Vials were placed at 29 °C for 24 h to induce rapid bacterial growth. After 24 h, we removed flies either for imaging or for CFU measurements (see **CFU measurements**). Flies that were kept for longitudinal imaging were placed in fresh vials at 25 °C with one male and flipped each day onto fresh food.

### CFU measurements

To quantify the bacterial load in flies, we used 3 flies per day kept in the same conditions as flies used for Bellymount imaging. Flies were kept on ice for 2-4 h and then washed with 70% ethanol three times and sterile PBS three times to remove external bacteria. We homogenized flies individually in 1.5 mL microcentrifuge tubes containing 100 µL sterile PBS using a motorized pestle. Next, we diluted the homogenate 10 times 1:5 in sterile PBS and spotted 3 µL onto MRS agar plates (1.5% agar) containing 10 µg/mL chloramphenicol. We counted colonies after growing for 48 h at 30 °C.

### Analysis of *L. plantarum* filling fraction

To quantify the bacterial density in the digestive tract following fly feeding, we measured the filling fractions of the crop and midgut. Images were analyzed using custom Matlab (R2018b) code. To calculate the filling fraction, the edge of the crop or the midgut was identified from the bacterial signal. The filling fraction was calculated as the pixel area occupied by cells ∼5 µm into the organ in the *z*-direction using a manually selected intensity threshold to capture signal from bacteria.

## Author Contributions

L.A.J.K., A.A.-D., J.L.M., W.B.L., K.C.H., and L.E.O. conceived the study; L.A.J.K., A.A.-D., Y.H.S., S.B., and J.L.M. performed the research; L.A.J.K., A.A.-D., K.C.H., and L.E.O. analyzed the data; and L.A.J.K., A.A.-D, K.C.H., and L.E.O. wrote the paper. All authors provided input on the manuscript.

## Acknowledgements

L.A.J.K was supported by 2T32GM00779038 and by Ruth L. Kirschstein Diversity Predoctoral Fellowship 1F31GM123736-01. A.A.-D. was supported by a Howard Hughes Medical Institute International Student Research Fellowship and a Stanford Bio-X Bowes Fellowship. This work was supported by NIH DP5OD017851 (to W.B.L.), the Allen Discovery Center at Stanford on Systems Modeling of Infection (to K.C.H), NIH RO1GM116000-01A1 (to L.E.O), and ACS RSG-17-167-01 DDC (to L.E.O.). K.C.H. is a Chan Zuckerberg Biohub Investigator. Confocal microscopy was performed at the Stanford Beckman Cell Sciences Imaging Facility (NIH S10RR02557401 and 1S10OD010580) and the Stanford Shriram Cell Sciences Imaging Facility funded by the Beckman Center and the Stanford School of Engineering, respectively. Initial CAD files were generated and printed by Jerome T. Geronimo at Stanford’s 3-Dimensional Printing & Rapid Prototyping Facility. We are grateful to have obtained *Drosophila* stocks from David Bilder, Carl Thummel, Yoshihiro Inoue, Pavel Tomancak, Joaquin de Navascues, the Kyoto Stock Center (DGRC), and the Bloomington Drosophila Stock Center (NIH P40OD018537). We thank Ihuicatl and Canciana for being highly cooperative Bellymount subjects; Linda Jaramillo Koyama, Jeffrey Wesley Carlson, Katharine Ng, and members of the Huang and O’Brien labs for helpful discussions; and Allen Spradling Irene Miguel-Aliaga, Minx Fuller, and Jon-Michael Knapp for valuable comments on the manuscript.

## Supplemental Figures 1-6

**Supplemental Figure 1:**
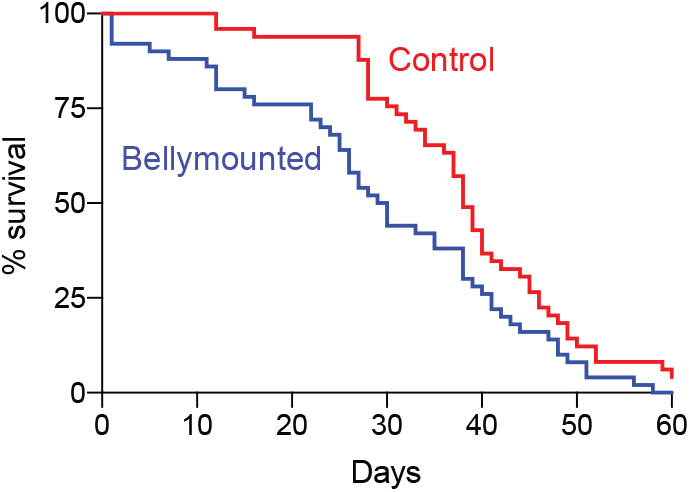
Longevity of animals after Bellymount. Adult females (4 days post-eclosion) were subjected to the Bellymount protocol (gluing and gentle compression) for 1 h and subsequently released. Lifespans of Bellymounted animals (*n*=50) were compared to a control cohort of age-and sex-matched animals that were neither glued nor compressed (*n*=49). One day after being released, 92% Bellymounted flies were alive. If successive applications of the Bellymount protocol each have the same effect on individual mortality, then population-level survival after three Bellymount protocols would be ∼78% (0.92^3^ = 0.78). Genotype: *ubi-his2avD::YFP*.

**Supplemental Figure 2:**
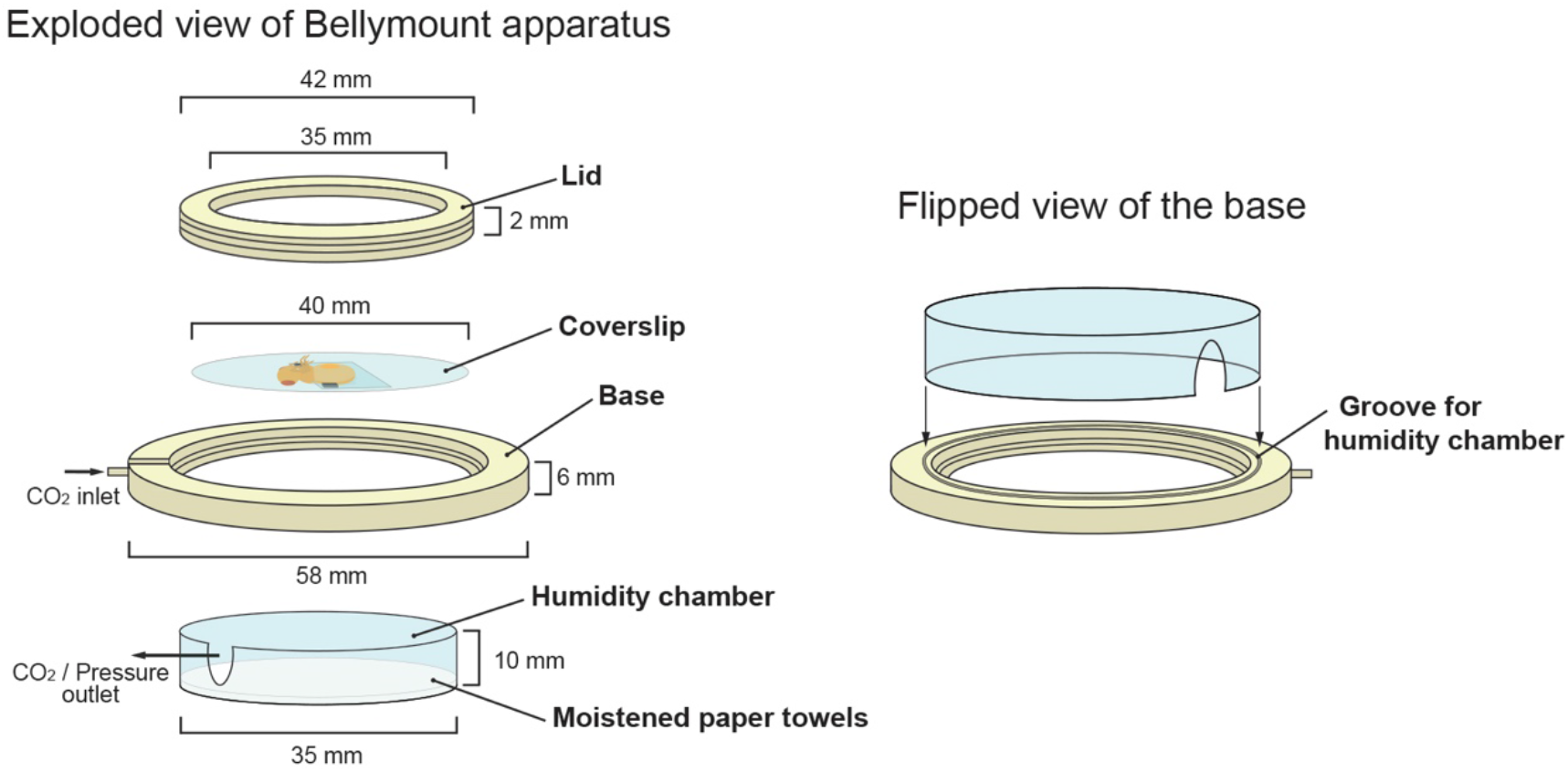
Exploded view of Bellymount apparatus. Isometric cartoon shows the arrangement of individual components in the assembled apparatus. A 40-mm coverslip with the glued, compressed fly is nested inside a custom base and secured by screwing on the lid. Supplemental Files 1 and 2 are CAD files for 3D printing of the lid and base. To prevent the fly from desiccating, a humidity chamber (35-mm petri dish containing H2O-soaked Kimwipes) attaches to a groove in the underside of the base.

**Supplemental Figure 3:**
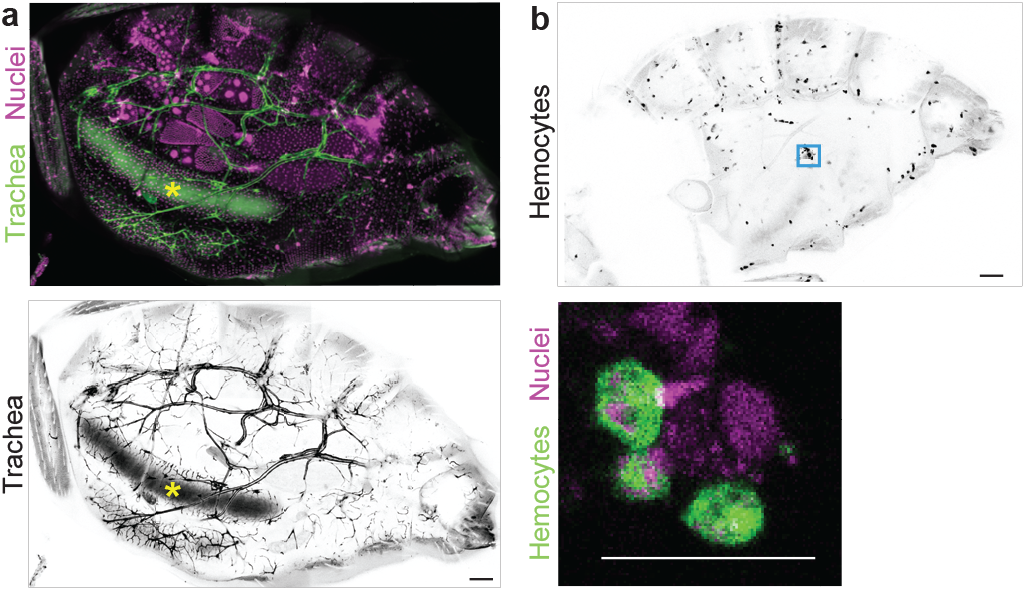
Abdominal trachea and hemocytes in Bellymounted animals. a, Abdominal tracheal network. Primary, secondary, and tertiary branches are visible (*btl>GFP*; green in top panel, inverted grayscale in bottom panel). Magenta (top) shows all nuclei (*ubi-his2av::mRFP*). Ingested food in the midgut lumen (yellow asterisk) has autofluorescence in the green channel. Scale bar, 100 µm. b, Abdominal hemocytes. *Top*, Whole-abdomen distribution (*hml>GFP*; inverted grayscale). *Bottom*, Zoom-in of boxed region in top panel shows single hemocytes (*hml>GFP*; green). Magenta shows all nuclei (*ubi-his2av::mRFP*). Scale bars, 100 µm (top) and 30 µm (bottom). All images are *z*-stack projections.

**Supplemental Figure 4:**
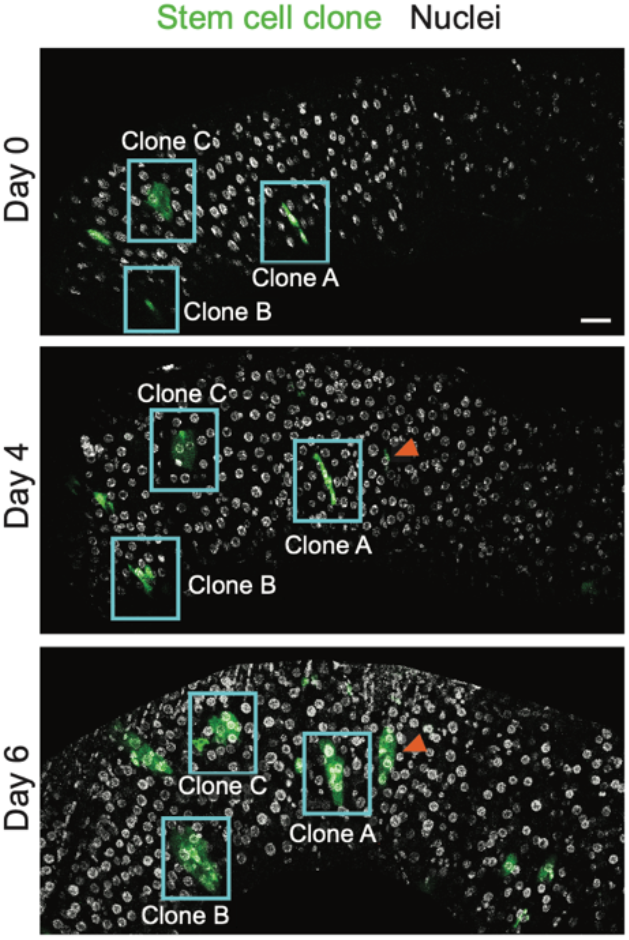
Persistent organ-level patterns enable re-identification of midgut clones. Wide-field view of Fly 3 midgut at each imaging timepoint (Fig. 1a). GFP-labeled stem cell clones were visible as green multicellular clusters. Blue boxes outline trackable clones that were analyzed in detail (Fig. 2d-l). Some spontaneous clones (orange arrowheads) developed over the duration of the experiment. Greyscale shows all nuclei (ubi-his2av::mRFP). Scale bar, 30 µm.

**Supplemental Figure 5:**
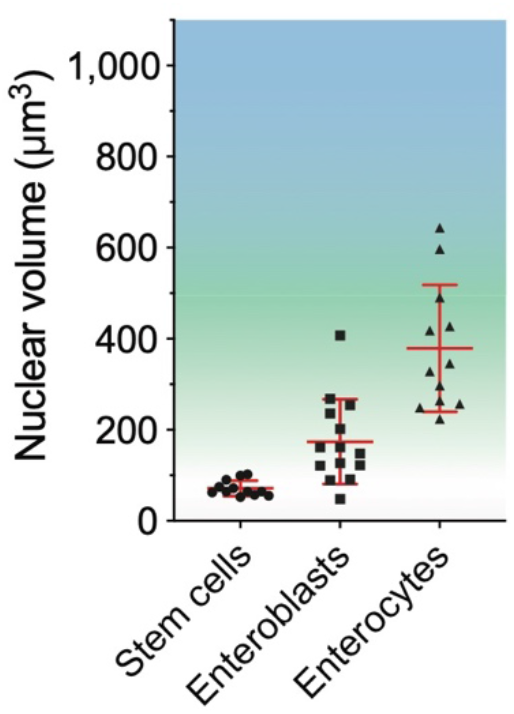
Enterocyte differentiation is characterized by increased nuclear volume. Nuclear volumes were measured from 3D reconstructions of fixed mid-guts that expressed cellular markers for successive stages of enterocyte differentiation: stem (and stem-like) cells, immature enteroblasts, and mature enterocytes. Mean ± S.D. nuclear volumes: stem cells (*n*=12), 71.8 ± 16.8 µm^3^; enteroblasts (*n*=14), 174.2 ± 93.1 µm^3^; enterocytes (*n*=12), 378.7 ± 139.3 µm^3^. Genotype: *esg>his2b::CFP, GBE-Su(H)-GFP:nls; ubi-his2av::mRFP*. Cell types were distinguished as follows: stem cells, RFP^+^ and CFP^+^; enter-oblasts, RFP^+^, CFP^+^, and GFP^+^; enterocytes, RFP^+^ and polyploid.

**Supplemental Figure 6:**
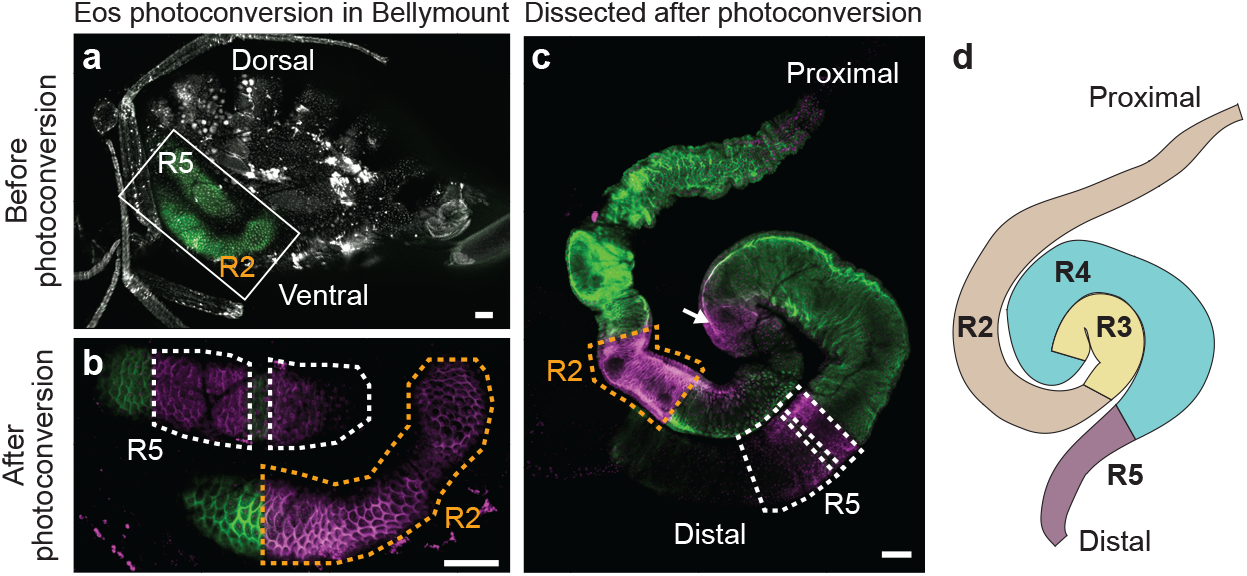
Proximal (R2) and distal (R5) midgut regions can be viewed in Bellymount. The photoconvertible protein Eos was used to determine which midgut regions are visible in Bellymounted flies. Female flies with midgut-specific expression of Eos (*mex>Eos*) were Bellymounted, and targeted areas were photoconverted by illumination with 405-nm wavelength laser light. Before photoconversion (a), Eos appeared as green fluorescence in the dorsal and ventral midgut loops (white box). After photo-conversion (b), Eos appeared red (magenta pseudocolor) in the laser-targeted areas (dotted outlines) and remained green in other areas. To determine which midgut regions were photoconverted, flies were removed from Bellymount and their midguts were dissected (c) and compared to the stereotyped anatomy of midgut regions R1-5 (d). This comparison showed that the photoconverted dorsal loop (white dotted outlines) was in R2 and the ventral loop (orange outline) was in R5. Weak photoconversion was also visible in R4 (arrow), likely because of its proximity to R2 *in situ.* The same pattern was observed in 100% (11/11) of photoconverted flies examined. All scale bars, 100 µm.

## Supplemental Movies

**Supplemental Movie 1: Release of Bellymounted fly after imaging.** Gentle prying with forceps removed a Bellymounted fly from Elmer’s Glue on the imaging coverslip. The fly was intact and immediately walked out of view.

**Supplemental Movie 2: Real-time nutrient ingestion, gastrointestinal transit, and intestinal peristalsis.** A Bellymounted fly was provided a cloth wick saturated with 5% sucrose water colored by Brilliant Blue FCF. Over the 45-min imaging session, the ingested liquid filled the crop and successive compartments of the midgut. Rapid peristaltic contractions of the midgut tube were visible.

**Supplemental Movie 3: Carbon dioxide (CO_2_) anesthesia inhibits tissue movement during confocal imaging.** Time-lapse confocal imaging of the midgut in a Bellymounted fly. Continuous imaging was performed while CO_2_ flow to the Bellymount apparatus was toggled on and off as indicated. With CO_2_ off, tissue movement caused cells to appear duplicated. With CO_2_ on, tissue movement was inhibited and no duplications were observed. Red marks nuclei in all tissues (*ubi-his2av::mRFP*). Green marks midgut stem cells and enteroblasts (*esgGal4>UAS-LifeAct-GFP*). Time intervals for CO_2_ off and on were 15 min and 20 min, respectively. Each movie frame is a *z*-stack projection. *Z*-stacks were captured at 5-min intervals, and each required ∼2 min to acquire 41 optical sections at intervals of 3 µm. Total movie time is 3.4 h.

**Supplemental Movie 4: Native arrangement of female abdominal organs at high resolution.** Animated *z*-stack of the tiled, whole-abdomen projection shown in Fig. 1f. Single optical sections through Bellymounted female were taken at 3-µm steps from the exterior cuticle to an interior depth of 54 µm. All nuclei are marked with His2av::mRFP (inverted grayscale). Scale is as indicated in Fig. 1f.

**Supplemental Movie 5: Volumetric image of developing egg chambers.** Three-dimensional reconstruction of egg chambers in the ovary of a Bellymounted female. Nascent oocytes and supporting nurse and follicles were readily identifiable. Labels indicate eggs at different developmental stages, the surrounding fat body, and adjacent cuticle. All nuclei are marked with His2av::mRFP (grayscale). Scale bar, 30 µm.

**Supplemental Movie 6: Volumetric image of the crop lumen of a fly two days after colonization with *L. plantarum*-mCherry.** 3D reconstruction of same field as in Fig. 2o showing the *Drosophila* cuticle and crop imaged with Bellymount. *L. plantarum-*mCherry cells were visible in clumps and lined the edge of the crop lumen. Circular yeast cells were visible in the crop lumen. The external cuticle and body cavity between the cuticle and crop were visible. Greyscale indicates *L. plantarum-*mCherry signal and autofluorescence from yeast cells and the external cuticle. Scale bar, 10 µm.

**Supplemental Movie 7: Volumetric image of the proximal midgut of a fly two days after colonization with *L. plantarum-*mCherry.** 3D reconstruction of the same field as in Fig. 2q taken with Bellymount. The proximal midgut showed clumps of *L. plantarum*-mCherry cells as well as free-floating bacteria within the lumen. Yellow dotted lines mark the edge of the midgut lumen. Greyscale indicates *L. plantarum*-mCherry signal. Scale bar, 15 µm.

**Supplemental Movie 8: Volumetric image of the distal midgut of a fly two days after colonization with *L. plantarum-*mCherry.** 3D reconstruction of the same field in Fig. 2q taken with Bellymount. By contrast with the proximal midgut (Supplemental Movie 7), the distal midgut showed only free-floating bacteria within the lumen. Yellow dotted lines mark the edge of the midgut lumen. Greyscale indicates *L. plantarum-*mCherry signal. Scale bar, 15 µm.

## References

1. He, Y. & Jasper, H. Studying aging in Drosophila. Methods 68, 129–133 (2014).

2. Losick, V. P., Morris, L. X., Fox, D. T. & Spradling, A. Drosophila stem cell niches: a decade of discovery suggests a unified view of stem cell regulation. Developmental Cell 21, 159–171 (2011).

3. Hou, S. X. & Singh, S. R. Stem-cell-based tumorigenesis in adult Drosophila. Curr. Top. Dev. Biol. 121, 311–337 (2017).

4. Douglas, A. E. The Drosophila model for microbiome research. Lab Anim 47, 157–164 (2018).

5. Ugur, B., Chen, K. & Bellen, H. J. Drosophila tools and assays for the study of human diseases. Dis Model Mech 9, 235–244 (2016).

6. Al, K. et al. Live imaging of a Drosophila melanogaster model of nephrolithiasis with probiotic suplementation using CO_2_ anesthesia and Micro-CT. J. Urol. 201, e22–e23 (2019).

7. Poinapen, D. et al. Micro-CT imaging of live insects using carbon dioxide gas-induced hypoxia as anesthetic with minimal impact on certain subsequent life history traits. BMC Zoology 2, 9 (2017).

8. Men, J. et al. Optogenetic cardiac control in Drosophila using red-light. in BRAIN JW3A.33 (Optical Society of America, 2018).

9. Fichelson, P. et al. Live-imaging of single stem cells within their niche reveals that a U3snoRNP component segregates asymmetrically and is required for self-renewal in Drosophila. Nature Cell Biology 11, 685–693 (2009).

10. Lenhart, K. F. & DiNardo, S. Somatic cell encystment promotes abscission in germline stem cells following a regulated block in cytokinesis. Developmental Cell 34, 192–205 (2015).

11. Morris, L. X. & Spradling, A. C. Long-term live imaging provides new insight into stem cell regulation and germline-soma coordination in the Drosophila ovary. Development 138, 2207–2215 (2011).

12. Martin, J. L. et al. Long-term live imaging of the Drosophila adult midgut reveals real-time dynamics of division, differentiation and loss. eLife 7, e36248 (2018).

13. Apidianakis, Y., Tamamouna, V. & Teloni, S. Intestinal Stem Cells: A decade of intensive research in Drosophila and the road ahead. Advances in Insect … 52, 139–178 (2017).

14. Hsu, Y.-C. & Fuchs, E. A family business: stem cell progeny join the niche to regulate homeostasis. Nat Rev Mol Cell Biol 13, 103–114 (2012).

15. Lee, T. & Luo, L. Mosaic analysis with a repressible cell marker for studies of gene function in neuronal morphogenesis. Neuron 22, 451–461 (1999).

16. Liang, J., Balachandra, S., Ngo, S. & O’Brien, L. E. Feedback regulation of steady-state epithelial turnover and organ size. Nature 548, 588–591 (2017).

17. de Navascués, J. et al. Drosophila midgut homeostasis involves neutral competition between symmetrically dividing intestinal stem cells. EMBO J 31, 2473–2485 (2012).

18. Ohlstein, B. & Spradling, A. The adult Drosophila posterior midgut is maintained by pluripotent stem cells. Nature 439, 470–474 (2006).

19. Klein, A. M. & Simons, B. D. Universal patterns of stem cell fate in cycling adult tissues. Development 138, 3103–3111 (2011).

20. Wong, C. N. A., Ng, P. & Douglas, A. E. Low-diversity bacterial community in the gut of the fruitfly Drosophila melanogaster. Environmental Microbiology 13, 1889–1900 (2011).

21. Tropini, C., Earle, K. A., Huang, K. C. & Sonnenburg, J. L. The gut microbiome: Connecting spatial organization to function. Cell Host Microbe 21, 433–442 (2017).

22. Willis, A. R. et al. Shigella-induced emergency granulopoiesis protects zebrafish larvae from secondary infection. mBio 9, e00933–18 (2018).

23. Obadia, B. et al. Probabilistic invasion underlies natural gut microbiome stability. Current Biology 27, 1999–2006.e8 (2017).

24. Stoffolano, J. G. & Haselton, A. T. The adult Dipteran crop: a unique and over-looked organ. Annu. Rev. Entomol. 58, 205–225 (2013).

25. Shanbhag, S. & Tripathi, S. Epithelial ultrastructure and cellular mechanisms of acid and base transport in the Drosophila midgut. J Exp Biol 212, 1731–1744 (2009).

26. Wiedenmann, J. et al. EosFP, a fluorescent marker protein with UV-inducible green-to-red fluorescence conversion. PNAS 101, 15905–15910 (2004).

27. O’Brien, L. E., Soliman, S. S., Li, X. & Bilder, D. Altered modes of stem cell division drive adaptive intestinal growth. Cell 147, 603–614 (2011).

